# Whole-brain calcium imaging during physiological vestibular stimulation in larval zebrafish

**DOI:** 10.1101/300350

**Authors:** Geoffrey Migault, Thomas Panier, Raphaël Candelier, Georges Debrégeas, Volker Bormuth

## Abstract

During in vivo functional imaging, animals are head-fixed and thus deprived from vestibular inputs, which severely hampers the design of naturalistic virtual environments. To overcome this limitation, we developed a miniaturized ultra-stable light-sheet microscope that can be dynamically rotated during imaging along with a head-restrained zebrafish larva. We demonstrate that this system enables whole-brain functional imaging at single-cell resolution under controlled vestibular stimulation. We recorded for the first time the dynamic whole-brain response of a vertebrate to physiological vestibular stimulation. This development largely expands the potential of virtual-reality systems to explore complex multisensory-motor integration in 3D.

---

During natural behaviors, motor action and sensory perception are intertwined in a feedback loop. Virtual reality provides a powerful method to investigate this sensory-motor coupling^8,18^. It allows to freely manipulate the relationship between sensory inputs and motor action in order to study sensory-motor integration processes, and this even in experimental contexts that do not exist in real world. Furthermore, virtual reality systems can evoke naturalistic behaviors in head-restrained animals making the brain accessible to simultaneous brain-activity monitoring, using calcium imaging or neurophysiological methods^12^. Such approaches have been successfully implemented in mice, drosophilae and zebrafish^10^. In the latter system, the entire brain was monitored in vivo using light-sheet imaging as the animal visually interacted with its environment^1,2,15,19–21^.

However, tethering the animal’s head inevitably disrupts vestibular feedback of self-motion and of body orientation relative to the gravitational axis, leading to sensory mismatch and altered neuronal processing. Compared to freely running assays, electrophysiological recordings in mice navigating in virtual 2D space show modified spatial coding by hippocampal place cells, reflecting the lack of vestibular feedback^4^. As fish navigate in 3D space, their vestibular system provides continuous information about postural orientation relative to gravity as well as about translational and rotational acceleration. Vestibular information is involved in multisensory driven locomotion initiation^9^, postural control and gaze stabilization^6,14^. Vestibular cues also enable larvae zebrafish to find the water surface, to inflate the swim bladder or to navigate to the ground when stressed. Vestibular deficient animals are not viable^16^.

Large-scale calcium imaging is currently incompatible with controlled vestibular stimulation making it impossible to integrate this sensory modality into existing virtual reality systems. The technical challenge arises as physiological stimulation of the vestibular system demands dynamic rotation and tilting as well as translation of the animal. If performed under a fixed microscope, this would constantly change the imaged brain section and thus the set of neurons under investigation making it difficult to monitor the activity of individual neurons over time.

Here we report on the development of a light-sheet microscope that can be rotated during whole-brain recording in behaving larval zebrafish. Rotating the microscope along with the head-restrained larvae enables us to stimulate the vestibular system in a controlled and physiological way while maintaining imaging conditions unchanged, with sub-micron precision (Fig. 1a). Imaging conditions and data analysis pipelines are the same as for other stimulation types as the animal remains static relative to the microscope. Existing virtual reality devices for other modalities integrate thus directly into our system.

**Figure 1:**
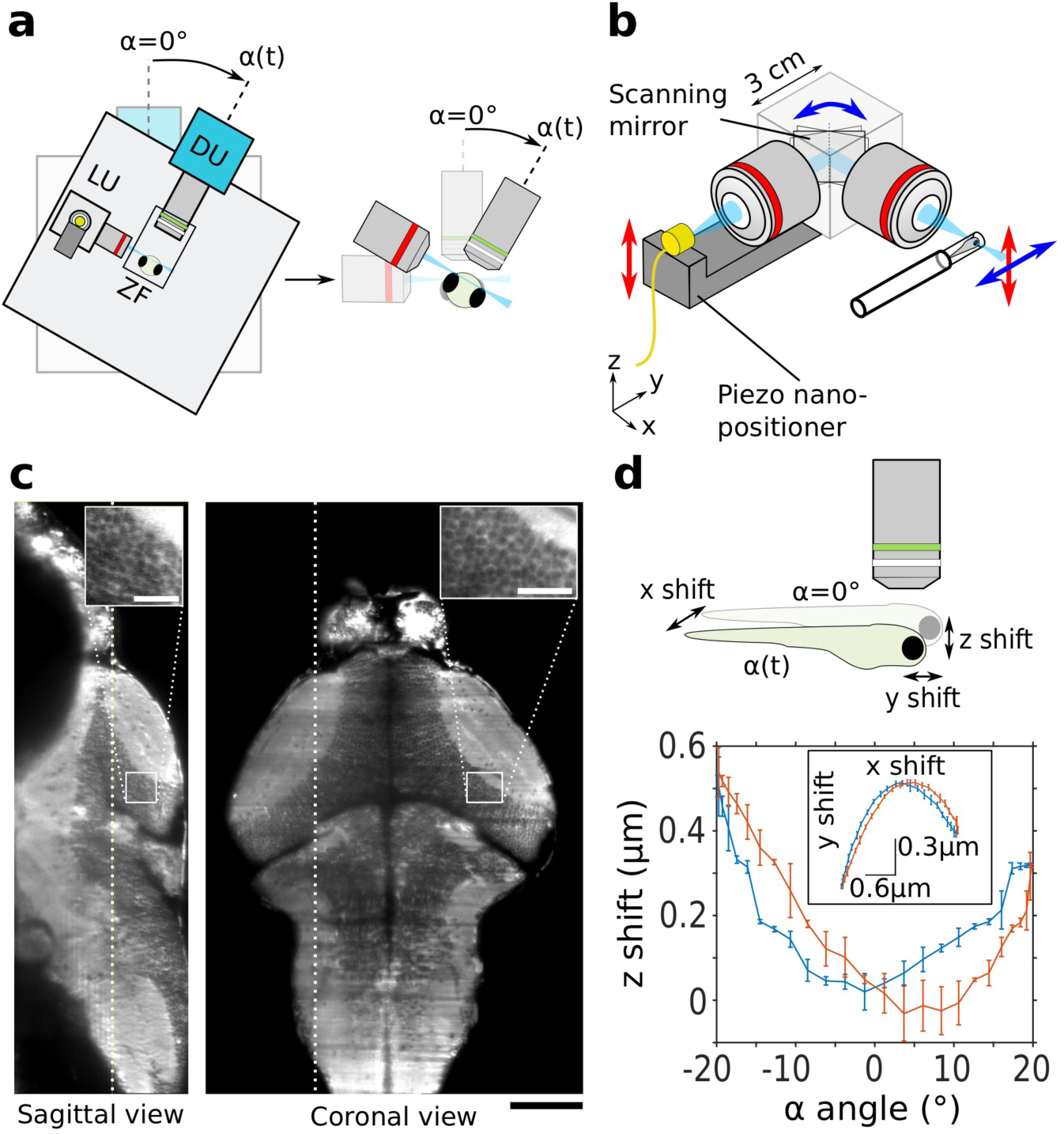
Experimental setup and performance characterization. (a) Schematic of the rotating microscope setup with the light-sheet unit (LU), the mounted zebrafish (ZF) and the detection unit (DU). The zoom shows that the fish rotates together with the microscope such that the light-sheet and the detection focal plane do not depend on the microscope rotation angle. Stimulation angles are counted positive in the direction indicated by the arrow. The fish looks onto the breadboard with its long body axis aligns onto the microscope rotation axis. (b) Schematic of the light-sheet forming unit. (c) Sagittal and coronal section of a volumetric whole-brain scan of a 6dpf old zebrafish larva with pan-neuronal GCaMP6s expression. Scale bar 100μm. Insets shows single-cell resolution. Scale bar 20μm. (d) Characterization of the scanned volume shift along the z-direction (see schematic) as function of the microscope rotation angle. Rotation direction from −20° to +20° in blue and reversed rotation in red. The inset shows the corresponding movements in the x-y plane.

Rotating the microscope while performing high-resolution neuronal recordings imposes stringent demands on the mechanical rigidity of the optical apparatus. Any mechanical distortion resulting from the changing gravitational forces during microscope rotation may indeed induce displacements of the imaging plane. Such displacements would transiently bring another set of neurons in focus, leading to detrimental artefacts in the fluorescence recordings. To overcome these challenges, we designed a novel miniaturized illumination unit for 3D digitally-scanned light-sheet microscopy. By reducing the optical components to the minimal necessary set, we were able to obtain a ten-fold reduction in size of the optical path compared to standard digital scan light-sheet microscopes. This compact design eliminates most mechanical distortion and makes the microscope insensitive to vibrations.

To produce the light-sheet, the output of an optical fiber is imaged into the sample using two low NA objectives and a scanning mirror (Fig. 1b). In this design, we omitted a relay system that is conventionally used to bring the pivoting point of the mirror onto the back focal plane of the illumination objective to ensure parallel beam deflection. However, we calculated that the resulting laser pointing variation across the ~mm^2^ field of view is less than 0.4° (see Supplementary Discussion and Fig. S1), and, as it occurs within the light-sheet plane, it has no impact on the imaging performance. The divergence of the beam at the fiber output further offers an adequate numerical aperture for micron-thick light-sheet without the need to introduce a beam-expander system. Noticeably, this configuration compensates for spherical aberrations as the laser passes through the two identical objectives in reversed direction. Z-scanning (Fig. 1b-c, Fig. S1b and Movie S1) is obtained by directly moving the fiber outlet using a piezo-crystal, which in turn displaces the light-sheet. This is possible as the telescope formed by the two objectives conjugates the light-sheet waist to the fiber output. Although the presence of the scanning mirror between the two objectives abolishes perfect telecentricity, the resulting pointing error is less than 0.2° and corresponds to a displacement of the light-sheet relative to the focal plane of less than 1.2 % of the light-sheet thickness. The unit dimensions are only 9×6×10cm (Fig. 1b and Fig. S1c-d) and the light-sheet can be aligned in a straightforward way by rotating and translating the entire unit. All optical elements - including the fluorescence detection unit, a behavior tracking unit and the sample holder - are mounted onto a breadboard (Fig. 1b) attached onto an ultra-stable high-load rotation stage (ALAR150SP, Aerotech, USA) which can accelerate the microscope with up to 10000°/s^2^ to an angular velocity of 60°/s in less than 10ms without any backslash

Our system provides high-resolution single-cell resolved volumetric brain scans (Fig. 1c and Movie S1). The high-quality light-sheet profile is consistent with the predictions of Gaussian optics with a half width at 1/e^2^ of 2.04 ± 0.02 μm (std) (Supplementary and Fig. S3). During sinusoidal microscope rotation over ± 20° of amplitude the scan volume moves by less than 500 nm in the z direction (Fig. 1d, Fig. S4 and Supplementary Method) which is only ~6 % of the typical soma diameter in zebrafish larvae (8um). The resulting noise in the fluorescence signal (ΔF/F) during calcium imaging sessions is less than 3% (std) over the entire range of rotational amplitude and acceleration (Fig. S5 and Supplementary Method). Even during microscope rotation, the imaging quality of our miniaturized setup in terms of signal-to-noise ratio and spatial resolution thus remains equivalent to state-of-the-art standard whole-brain functional light-sheet imaging.

In the following we illustrate how this system enables whole-brain functional imaging at single-cell resolution under controlled vestibular stimulation. We mapped the whole-brain response to dynamic vestibular stimulation of larval zebrafish expressing the calcium reporter GCaMP6s^19^ pan-neuronally. 6-day old larvae, partially embedded in agarose so that they can freely move their eyes, where submitted to sinusoidal rolling oscillations about their rostro-caudal axis at 0.2 Hz frequency and ± 10° amplitude. The fish responded to this rolling stimulation with anti-phasic compensatory eye movements of similar amplitude such as to maintain clear vision (Fig. 2a). Simultaneously recorded brain-wide neuronal activity is shown in Movie S2. The measured fluorescent dynamics is rich, correlated with the stimulation, localized to specific brain regions and antisymmetric with respect to the two brain hemispheres. Activity in the optic tectum, induced by the motion of the eyes that creates retinal shift, is abolished in paralyzed preparations (Fig. S7).

**Figure 2:**
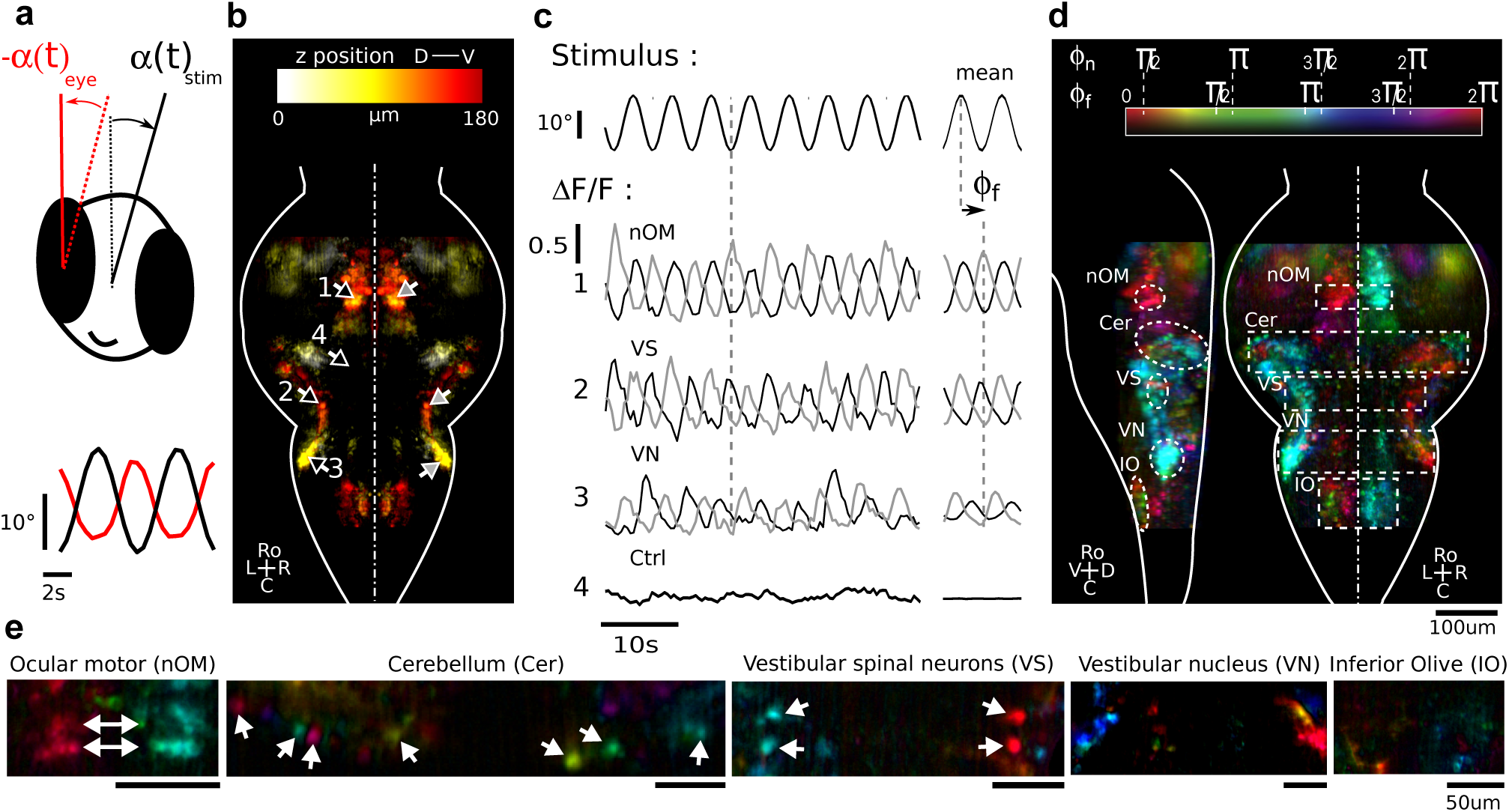
Functional brain-wide response to sinusoidal vestibular rolling stimulation. (a) Schematic illustrates the rotation of the fish when the microscope rotates and the evoked compensatory eyes movements. The fish looks into the plane and angles are counted positive in clockwise direction. Sinusoidal rolling stimulation (0.2Hz, ±10°, black) and associated anti-phasic compensatory eye movements (red). (b) Projection of the neuronal response map. The color indicates the z-location in the brain from dorsal to ventral. This map was obtained by averaging over 3 fish, after morphological registration and symmetrization of the individual maps by keeping only voxels with a counterpart on the contralateral brain hemisphere (see Supplementary). (c) Example fluorescent traces of neurons symmetrically located with respect to the midline and indicated by the arrows in (b). Time traces corresponding to neurons on the left (right) side are black (gray). (d) Lateral (left to midline) and dorsal projection views of the phase maps of the neuronal signal. The color indicates the phase-shift ϕ_f_ shown in (b) of the fluorescence signal with respect to the stimulation signal. Φ_n_ indicates the corresponding colors after correction by a uniform value to account for the GCaMP6s response function (see Supplementary). (e) Coronal sections of the phase-map of regions marked in (d) for a single fish.

We computed the brain-wide vestibular response map to rolling (average of N=3 fish) (Fig. 2b) by identifying all brain voxels whose fluorescence displays a significant oscillatory signal at the stimulation frequency (see Supplementary). The identified network is highly stereotyped with a bilaterally symmetric organization. Fluorescence traces of selected brain regions are shown in Figure 2c revealing different phases relative to the stimulation signal. The corresponding phase map (Fig. 2d, Fig. S6 and Movie S3) reveals two anti-phasically active subnetworks (red and cyan), comprising neurons in the vestibular nucleus, the cerebellum, the oculomotor nuclei, the nucleus of the medial longitudinal fasciculus and in hindbrain regions consistent with known circuit diagrams^3,6^. Furthermore, we observed neuronal populations whose activity is ~90° phase shifted relative to these two networks as e.g. in the vestibular nucleus (blue and yellow). Noticeably, we found high phase variability among the cerebellar Purkinje cells population as previously reported by electrophysiological recordings ^17^ (Fig. 2e).

Beyond this continuous stimulation protocol, the setup allows one to examine the whole-brain response to vestibular step-stimulation in the form of fast transient angular changes followed by periods of fixed angular position (Fig. 3 and Movie S4). These experiments revealed three classes of individual response types: (i) strong tonic fluorescent responses (tuned to the angular position of the fish) e.g. in the vestibular nucleus or the vestibular spinal neurons, (ii) mostly phasic responses (tuned to the angular velocity of the fish) e.g. in the tectum, and (iii) linear combinations of both, e.g. in the ocular motor neurons.

**Figure 3:**
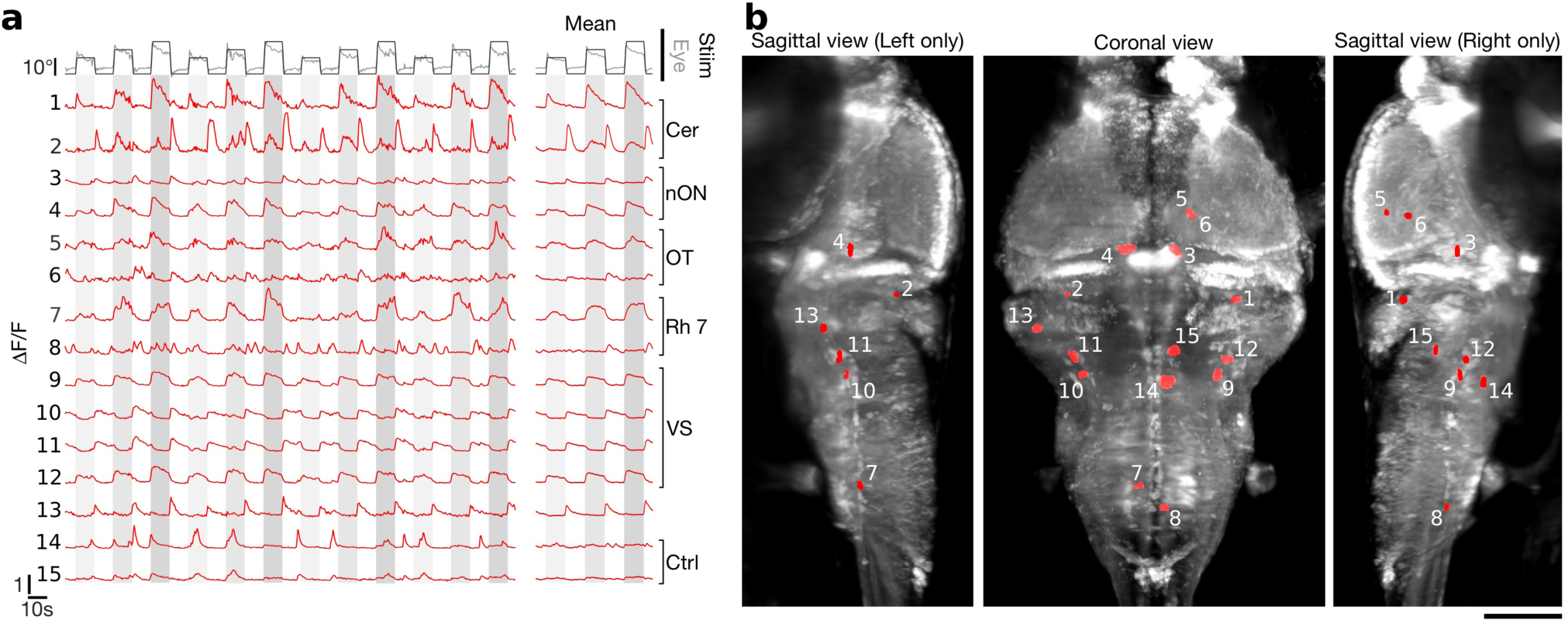
Functional brain-wide response to vestibular rolling step stimulation. (a) Time trace of the imposed vestibular stimulus (microscope rotation) with peak angular speed of 30°/s (black), compensatory eye movements (gray), fluorescent responses (red) measured in different brain regions indicated by numbers in (b). Averages are calculated over 10 stimulation cycles. (b) Three brain projections showing the location of the region-of-interests (red) from which the fluorescent time traces shown in (a) were extracted. The sagittal left (right) view is a projection from left (right) to the midline. The coronal view is a projection from dorsal to ventral. Scale bar 100μm

An alternative approach to deliver vestibular cues to head-fixed larvae has recently been demonstrated, which consists of directly moving utricular otoliths using optical tweezers^11^. Although such manipulations do evoke small compensatory motor behaviors^11^, it is difficult to calibrate, which precludes the delivering of well-controlled and reproducible physiological stimuli. A complementary avenue to virtual reality systems may also be provided by functional recording in freely swimming fish, as was recently proposed^7,13^. Beyond the lower SNR offered by this method, it lacks the possibility to disentangle vestibular from motor variables and is restricted to 2D trajectories.

With our rotating light-sheet microscope we successfully combine physiological dynamic vestibular stimulation and behavioral monitoring with single-cell resolved and high signal-to-noise ratio whole-brain calcium imaging. To achieve the exquisite mechanical stability required for such recordings, we designed a miniaturized digital scan light-sheet unit. This unit is of general interest because of its reduced footprint, which allows to transform virtually any microscope into a digital scan light-sheet microscope with high image quality. Due to the small number of optical components, it is relatively inexpensive, simple to build and straightforward to align. Further stimulation degrees of freedom can be implemented in a straight forward manner with commercially available motorized stages. Finally, the system is compatible with optical enhancements such as two-photon light-sheet function imaging^21^, line-confocal^5^ and imaging with multiple light-sheets^19^. With this platform, we recorded for the first time the dynamic whole-brain response of a vertebrate to vestibular stimulation. This development largely expands the potential of virtual-reality systems to explore complex multisensory-motor integration in 3D.

## Acknowledgements

We thank the IBPS fish facility staff, and in particular Alex Bois and Stéphane Tronche for their invaluable help. We are also grateful to Carounagarane Dore for his important contribution to the realization of the mechanical parts of the light-sheet microscope. This project has received funding from the European Commission under the Horizon 2020 research and innovation program Grant agreement (ERC starting grant No J17E298), by the ATIP-Avenir program from CNRS and Inserm, and by the ANR-16-CE16-0017. Geoffrey Migault was funded by a PhD fellowship from the Doctoral School in Physics, Ile de France (EDPIF).

